# Changes in the milk metabolome of the giant panda (*Ailuropoda melanoleuca*) with time after birth – Three phases in early lactation and progressive individual differences

**DOI:** 10.1101/029363

**Authors:** Tong Zhang, Rong Zhang, Liang Zhang, Zhihe Zhang, Rong Hou, Hairui Wang, I. Kati Loeffler, David G. Watson, Malcolm W. Kennedy

## Abstract

Ursids (bears) in general, and giant pandas in particular, are highly altricial at birth. The components of bear milks and their changes with time may be uniquely adapted to nourish relatively immature neonates, protect them from pathogens, and support the maturation of neonatal digestive physiology. Serial milk samples collected from three giant pandas in early lactation were subjected to untargeted metabolite profiling and multivariate analysis. Changes in milk metabolites with time after birth were analysed by Principal Component Analysis, Hierarchical Cluster Analysis and further supported by Orthogonal Partial Least Square-Discriminant Analysis, revealing three phases of milk maturation: days 1-6 (Phase 1), days 7-20 (Phase 2), and beyond day 20 (Phase 3). While the compositions of Phase 1 milks were essentially indistinguishable among individuals, divergences emerged during the second week of lactation. OPLS regression analysis positioned against the growth rate of one cub tentatively inferred a correlation with changes in the abundance of a trisaccharide, isoglobotriose, previously observed to be a major oligosaccharide in ursid milks. Three artificial milk formulae used to feed giant panda cubs were also analysed, and were found to differ markedly in component content from natural panda milk. These findings have implications for the dependence of the ontogeny of all species of bears, and potentially other members of the Carnivora and beyond, on the complexity and sequential changes in maternal provision of micrometabolites in the immediate period after birth.

---

**Abbreviated title** – Early lactation changes in giant panda milk.

## Introduction

The chemical complexity of milk required to support neonatal mammalian development is only partially understood. Beyond the basic functions of nutrition, milk provides immune protection and helps to establish a healthy gut microbiota in young mammals (1-3). Moreover, the composition of milks must meet the needs of nursing young at changing developmental stages. The ontogeny of ursid milk is particularly interesting because of the highly altricial nature of bear cubs and the long period of maternal dependence in ursidae. Giant panda cubs in particular are the most altricial of any non-marsupial mammal (4).

While marsupials produce the developmentally most immature neonates of all mammals, Ursidae follow close behind despite a eutherian physiology, taxonomic classification among Carnivora and small litter sizes. The evolutionary significance of this ursine characteristic remains unexplained. It may be an adaptation to the frugality with which bear sows manage their metabolic resources during hibernation, which is when cubs are born to those bear species that hibernate. Lactation transfers energy in the form of fats to the neonate, which is metabolically much less costly to the mother than the transplacental transfer of glucose to support the foetus (5-7).

The production of altricial neonates necessitates a particularly dynamic early lactation phase to meet the developmental needs of nursing young. We define the early lactation phase as that during which the neonate is dependent on maternal milk for provision of immune factors and maturation of the gut microbiota, as well as to meet changing physiological and metabolic needs. While the characteristics of early lactation have been studied extensively in marsupials (8, 9), studies of bear milks have heretofore been confined to analysis of basic nutritional components (10-16) or identification of oligosaccharides from single samples (17-22). We hypothesized that the early lactation phase of bears – specifically of giant pandas – is prolonged in comparison to that of other eutherian mammals, and that this phase presents discrete and dynamic changes in the metabolome profile.

We chose to work with giant pandas for two reasons. First, this species produces the most altricial young even of bears, with the body weight ratio between mother and new-born approximately 1000:1 and a markedly underdeveloped lymphoid system (4). Second, the intrusive methods of captive giant panda husbandry in China allow the collection of serial milk samples from animals trained to accept this interference. This type of sample collection is unavailable for other species of bears. As such, the panda situation presented a unique opportunity to investigate ursine lactation.

Urashima et al. carried out a series of studies to characterise oligosaccharides in single milk samples collected from various bear species, including the giant panda (17-22). In these studies, oligosaccharides present at high levels were isolated by anion exchange chromatography and identified by nuclear magnetic resonance spectroscopy. Their abundances were approximated by the published chromatogram in each study. While lactose is the dominant saccharide in, for example, human and bovine milks, these studies revealed that isoglobotriose dominates in the milks of bears. Studies of polar bear milk further indicated that the abundance of isoglobotriose changes from colostrum to mature milk (19). The analysis of giant panda milk in previous work was based on a single sample, and only the major oligosaccharides were isolated and identified (23). The present study sought to expand this with analysis of serial samples taken from three individuals in order to better understand the dramatic changes known to occur in the milks of bears during early lactation.

Metabolomics methodologies have been increasingly applied to milk research (24-34). These are designed for analysis of small molecules (MW <1500 kDa) in biological samples. These methods are associated with multivariate analysis, and make possible the comparison of metabolome fingerprints from multiple samples under different biological conditions. This is not possible with conventional analytical strategies. Mass spectrometry (MS) coupled with gas or liquid chromatography (GC or LC) is more widely applied than nuclear magnetic resonance spectroscopy in untargeted metabolite profiling studies because of its high sensitivity and ability to identify individual compounds by chromatographic retention time and tandem mass spectrometry (MS/MS) fragmentation patterns. These techniques have identified compounds that differentiate milks according to methods of production and processing (24). They also differentiate milks from individual animals according to lactation interval, and correlate metabolite profiles with various milk traits (24, 30). To date, metabolomics has been used to study the milks of humans and domesticated dairy species [12-22].

In the study presented here, we carried out untargeted metabolite profiling of milk samples from captive giant pandas, using a Hydrophilic Interaction Liquid Chromatography - High Resolution Mass Spectrometry (HILIC-HRMS) method. By using multivariate analysis, Principal Components Analysis (PCA) and Hierarchical Cluster Analysis (HCA), the changes in small molecule classes were mapped from serial samples collected from three pandas during early lactation. Further characterisation was conducted using MS/MS for key chemical components discovered from the multivariate analysis. We also trialled the application of Orthogonal Partial Least Square (OPLS) regression analysis to identify particular nutrient components correlated with the growth rates of cubs, using data from one giant panda cub for which we had a sufficient frequency of samples.

Three types of artificial milk formulae were also analysed. These products are used to supplement maternal milk while cubs are nursing, and to replace maternal milk once cubs have been removed from their mothers. Giant panda cubs are removed from their mothers in December, at 3-6 months of age, as part of standard husbandry practices in China’s giant panda breeding facilities, in order that the mother comes into oestrus again the following spring. Naturally, cubs stay with their mothers for 2.5 – 3.5 years, during which time the mother does not reproduce.

This study shows distinct, time-dependent changes in giant panda milk during early lactation that we discriminate into three phases. Further, our methods resolve differences among individual giant panda mothers in their milk metabolomes that develop with time in the course of lactation.

## Materials and methods

### Milk sample collection

Fifty-five milk samples were collected from three captive giant pandas during the 2012 parturient season at the Chengdu Research Base of Giant Panda Breeding, Chengdu, Sichuan Province, P.R. China. Sample collection and maternal information are included in Supporting Information Table S1. The growth rates of the cubs from these females, from birth over the first sixty days, are shown in Supporting Information Fig S1.

Samples were collected manually by giant panda keepers from females who had been trained previously to allow this manipulation without physical or chemical restraint. Sample volumes were between 1.5 and 4.5 ml, which were considered small enough to ensure that the milk source for the cub was not compromised, and to minimise any potential stress to the mother. Cubs were nursed by their mothers and supplemented daily with artificial milk formula, per standard husbandry practice at this facility.

Milk samples were frozen within two hours of collection and stored at -80°C, then transported frozen to the UK laboratory, after which they were stored at or below -20°C until analysis. Powdered human infant formula (Enfamil^®^, Mead Johnson, Indiana, USA) and puppy milk replacer (Esbilac^®^, PetAg, Illinois, USA) were obtained from commercial sources. These formulations are currently used directly or as a base for milk substitutes for giant panda cubs. A panda milk replacer (henceforth "Pandamilk"), formulated for research use, was obtained from the manufacturer (Morinyu Sunworld Co., Tokyo, Japan).

### Chemicals

High-performance Liquid Chromatography (HPLC) grade acetonitrile was obtained from Fisher Scientific, UK. Ammonium carbonate and ammonium hydroxide solution (28-30%) were purchased from Sigma-Aldrich, UK. HPLC grade water was produced by a Direct-Q3 Ultrapure Water System from Millipore, UK. The mixtures of metabolite authentic standards were prepared as previously described (35).

### Preparation of milk samples

The milk samples were thawed at room temperature, and 50μl of each were added to 200μl of methanol/acetonitrile 1:1 (v/v). The solution was mixed and placed in an ultra-sonic bath for 20 seconds. The emulsion was centrifuged for 10 minutes at 15,000 rpm at 4°C (Eppendorf 5424 R, maximum RCF = 21.130g). The supernatant was transferred to an HPLC vial for Liquid Chromatography-Mass Spectrometry (LC-MS) analysis. The three powdered milk formulae were prepared according to manufacturers’ instructions, and the extraction procedures were as above.

### HILIC-HRMS and multiple tandem HRMS analysis and data processing

Sample analysis was carried out on an Accela 600 HPLC system combined with an Exactive (Orbitrap) mass spectrometer (Thermo Fisher Scientific, UK). An aliquot of each sample solution (10μL) was injected onto a ZIC-pHILIC column (150 × 4.6mm, 5μm; HiChrom, Reading UK) with mobile phase A: 20mM ammonium carbonate in HPLC grade water (pH 9.2), and B: HPLC grade acetonitrile. The LC and the MS conditions were as described previously (36, 37). Samples were submitted in random order for LC-MS analysis, and the quality control sample (YY-19day) was injected at the beginning, middle, and end of the experiment to monitor the stability of the instrumentation. Peak extraction and alignment were calculated by integration of the area under the curve, using MZMine 2.10 software, as previously described (37). Resulting data were screened with an in-house macro (written in Microsoft Excel 2010) for removing the system background signals (cut-off for maximum signal intensity was 20 times control (blank)), and the remaining data were exported for multivariate analysis. Putative identification was also conducted in the Excel macro by searching the accurate mass against our in-house database (36, 37). Double or triple MS fragmentation of the key components discovered in the multivariate analysis was carried out with Collision Induced Dissociation at 35 V using a Surveyor HPLC system combined with a LTQ-Orbitrap mass spectrometer (Thermo Fisher Scientific UK). The MS/MS spectra were compared with published data or interpreted for further identification (Table S2).

The non-proton adducts and complex ions identified by MZMine 2.10 were confirmed by manually checking the raw LC-HRMS data. The importance of manual review of raw data is illustrated by the high occurrence (40%) of erroneous selection of LC-HRMS features by this type of processing. The remaining features were either confirmed by authentic chemical standards, or described by elemental composition analysis (accurate mass and isotope score) and multiple tandem MS fragmentation procedures by which their chemical classifications could be deduced.

### Multivariate analysis

SIMCA version 13.0 (Umetrics, Umeå, Sweden) was used for multivariate analysis including PCA, HCA, Orthogonal Partial Least Squares Discriminant Analysis (OPLS-DA) and OPLS. The data were centred, and unit variance scaled for PCA and HCA but Pareto scaled for OPLS-DA and OPLS in order to generate an S-plot for visualisation of the components with significant influence in the dataset. The rate of increase in cub body weight was used as the single Y-variable in OPLS. It was calculated as the percentage increase relative to the previous day’s weight (Fig S1).

### Ethics statements

Milk sampling from giant panda mothers was carried out under ethical approval from the University of Glasgow’s College of Medical, Veterinary & Life Sciences Ethics Committee, and the Chengdu Research Base of Giant Panda Breeding where the animals are held. All the giant panda mothers and their cubs were captive bred and maintained. Giant pandas are classified as endangered animals by the International Union for the Conservation of Nature (IUCN Red List 3.1; http://www.iucnredlist.org/details/712/0). International transfer of milk samples from captive giant pandas in China was covered by the CITES convention through permits issued by both donor and recipient countries. Milk samples were collected from conscious, unrestrained animals trained to allow milk sampling during routine health checks and veterinary inspection, or when considered necessary to provide milk for orphan or abandoned cubs, or for research purposes. Handling of cubs for health, normal growth, and veterinary checks is carried out daily at the Chengdu facility. A permit was obtained from the Scottish Executive for the importation of the milks sample as veterinary checked animal products to Scotland.

## Results and Discussion

### Slow progressive change in milk composition and increasing individuality with time after birth

After removing system interference, 2821 and 1945 liquid chromatography-high resolution mass spectrometry (LC-HRMS) features (pairs of mass to charge ratio (m/z) and retention time) were generated in electrospray ionisation negative and positive mode, respectively, and 368 features were detected in both modes. Of the generated features, 2199 compounds were characterised on the basis of accurate mass in our in-house database; 93 of these were further identified by matching their retention times with authentic standards (Table S3). The processed data were imported into SIMCA 13, and PCA was performed with mean centring and unit variance scaling. Eight principal components were generated with good model fitness and predictability (R^2^X (cum) = 61% and Q^2^ (cum) = 41.2%) for such a complex data matrix.

One of the aims of this study was to monitor the alteration of milk composition during the early stage of lactation and among individual giant pandas. To delineate these patterns, HCA was used to generate the dendrogram shown in Fig 1A from the PCA model. High repeatability of the LC-HRMS signals from three injections of the quality control sample (YY-19d) is reflected by the closeness of their locations in the HCA dendrogram and in the PCA score plot (the red circle in Fig 1A and 1B). This indicates excellent stability of our LC-HRMS instrumentation throughout the experiment.

**Fig 1.**
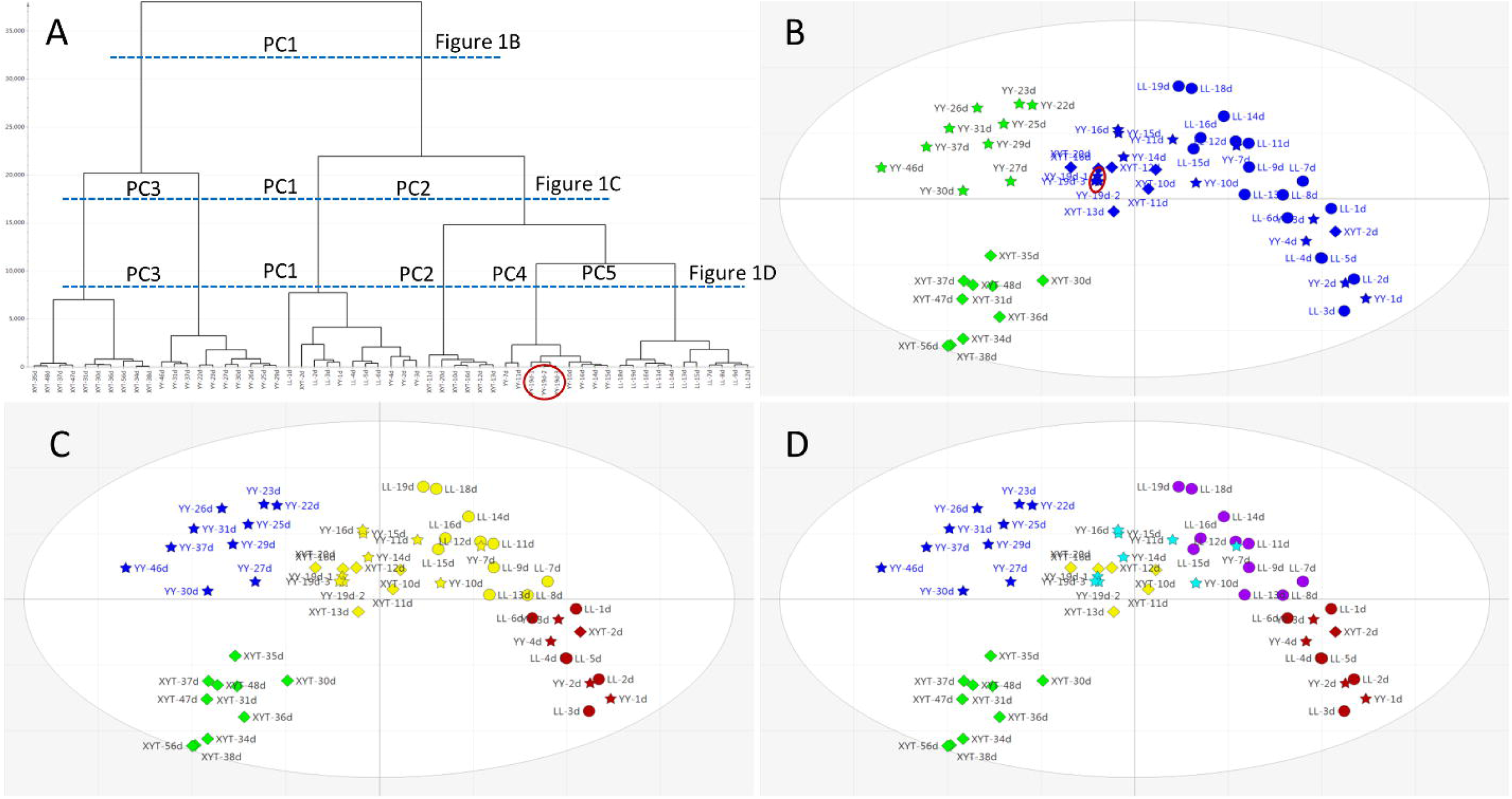
Progressive changes in milk composition and increasing individuality with time. Hierarchical cluster analysis (HCA) dendrogram (A) and principal components analysis (PCA) score plots (B-D) of milk samples collected serially from three giant pandas; Li Li (LL), circles; Yuan Yuan (YY), stars; Xiao Ya Tou (XYT), diamonds. Multivariate analysis using SIMCA (see Materials and Methods) allows horizontal sectioning of the dendrogram in (A) at different levels of clustering as indicated by the horizontal dotted lines. This yields progressively more detailed PCA score plots that first reveal similarities in metabolomes with time in all three individual giant pandas (as in (B), only two main coloured classifications), and then progressive disparities between the individuals when analysed at deeper resolutions (as in (C) and (D), increasing number of coloured classes). The data representing quality control repetitions (YY-19d-1, 2 and 3) are circled in red and indicate good repeatability. Score plots of PCA (B, C and D): x-axis PC1=19.4% and y-axis PC2=9.95%.

Observations in the PCA score plot were classified at points that corresponded to horizontal intersections of the dendrogram at certain levels. This revealed more principal components in the single, 2-dimensional (PC1 vs. PC2) PCA score plot. Figs 1A and 1B illustrate the first principal component separation of milk compounds at about 20 days postpartum, implying that there might be a significant alteration of the milk either in the components or in their relative abundance before and following that point. The early-phase group (the blue group in Fig 1B) was further separated by the second principal component into phases from days 1 to 6 (Fig 1C, red), and days 7 to 20 (Fig 1C, yellow). The later-phase (green) group in Fig 1B was divided into two groups by the third principal component, which distinguished individual animals (YY and XYT), rather than a further chronological separation (Fig 1C, blue and green.) The large gap between the blue and green groups in Fig 1C indicates the magnitude of difference between the milks of the two giant pandas after three weeks of lactation. Introduction of the fourth and the fifth principal components resulted in the pattern shown in Fig 1D, in which the 7-20-day (yellow) group in Fig 1C delineated components of the three individual pandas (purple, turquoise, yellow). Further classifications based on the remaining principal components are not shown here because of their lower impact on the PCA model.

The observations shown in Fig 1 indicate a time-dependent distribution of milk components, and further revealed unique profiles of individual animals. In the first 6 days of lactation, the component differences among the three pandas were relatively small, suggesting commonality in the composition of early milk among individual mothers. Differences subsequently became apparent with duration of lactation. On the basis of these observations, these giant panda milks may be classified into three phases: days 1-6 (Phase 1), days 7-20 (Phase 2) and following day 21 (Phase 3).

### Milk components that differ among the three phases of early lactation

In order to describe the relative abundance of components in Phase 1 milk samples, OPLS-DA was performed with Pareto scaling of compounds found in milks up to and including six days postpartum, and from seven days onward (Fig 2). The model (R^2^X (cum) = 89.9% and Q^2^ (cum) = 80.6%) was validated to high confidence by the permutation test (Figs S2.1 and S2.2). A clear separation was observed in the score plot from which the S-plot in Fig 2 was generated to visualise distinguishing components. Based on intensity and reliability, the 50 most abundant LC-HRMS features in pre-7 day samples, and the 20 most abundant features in post-7 day samples were highlighted in red and blue, respectively, in the S-plot. These 70 components are listed in Table 1.

**Fig 2.**
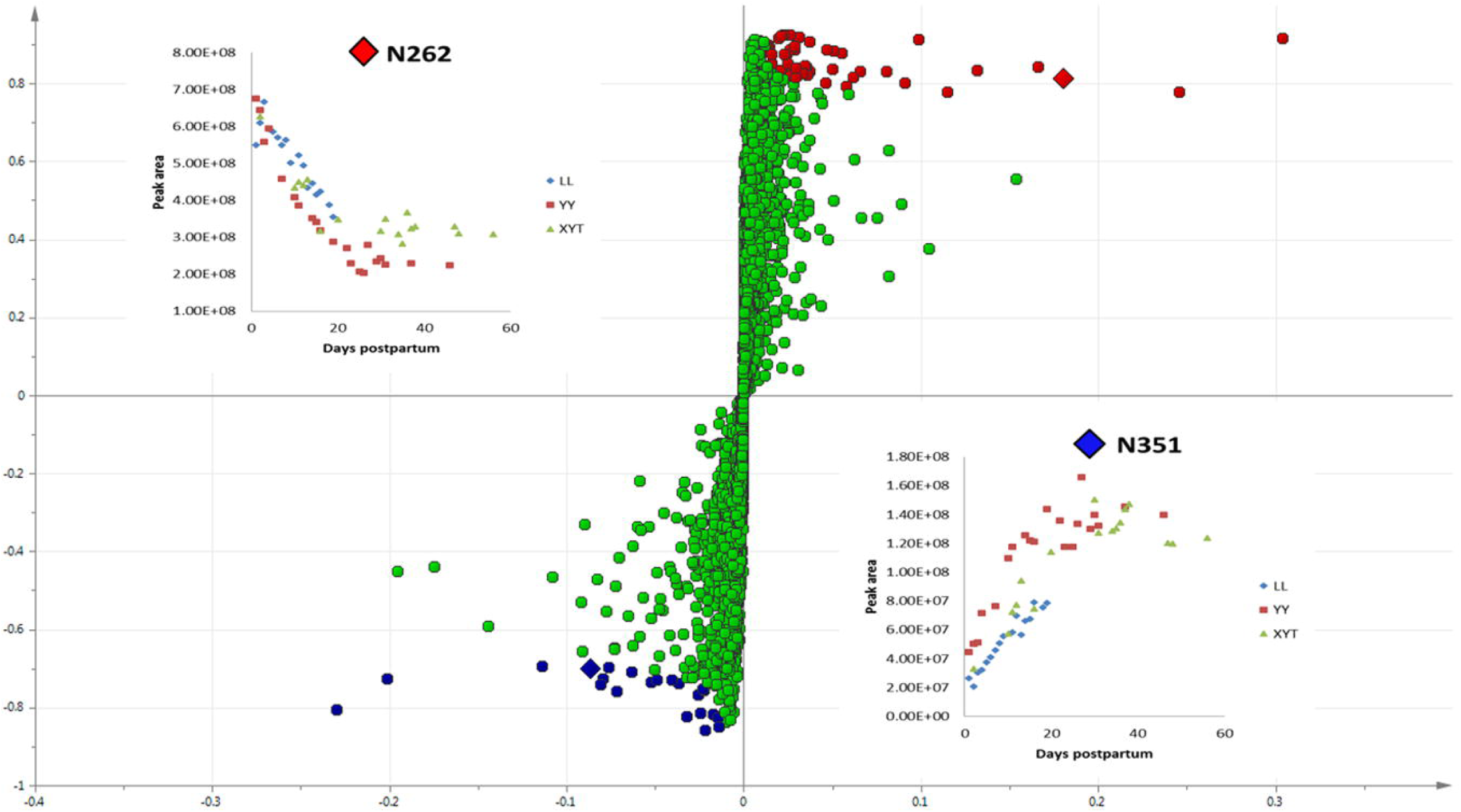
Changes in milk metabolome delineating discrete phases in early lactation. S-plot of orthogonal partial least squares discriminant analysis (OPLS-DA) of 55 giant panda milk samples collected serially after birth from three giant pandas. In an S-plot, the x variable is the relative magnitude of a variable, and the y variable is the variable confidence/reliability. So, data points falling in the upper right or lower left corners of the plot represent those features that are least likely to be the result of spurious correlations. Peaks with low magnitude/intensity falling in the centre of the plot near 0 are close to the noise level and exhibit high risks for spurious correlations. The 50 most abundant compounds in the early phase (before day 7; Phase 1) and the 20 most abundant in later milk (after day 7; Phases 2 and 3, cumulative) are highlighted in red and blue, respectively. The inset graphs describe the time-dependent changes in abundance of two oligosaccharide isomers N262 and N351 (identified as diamonds in the S-plot), subsequently identified as 3’-sialyllactose and 6’-sialyllactose, respectively.

**Table 1.**
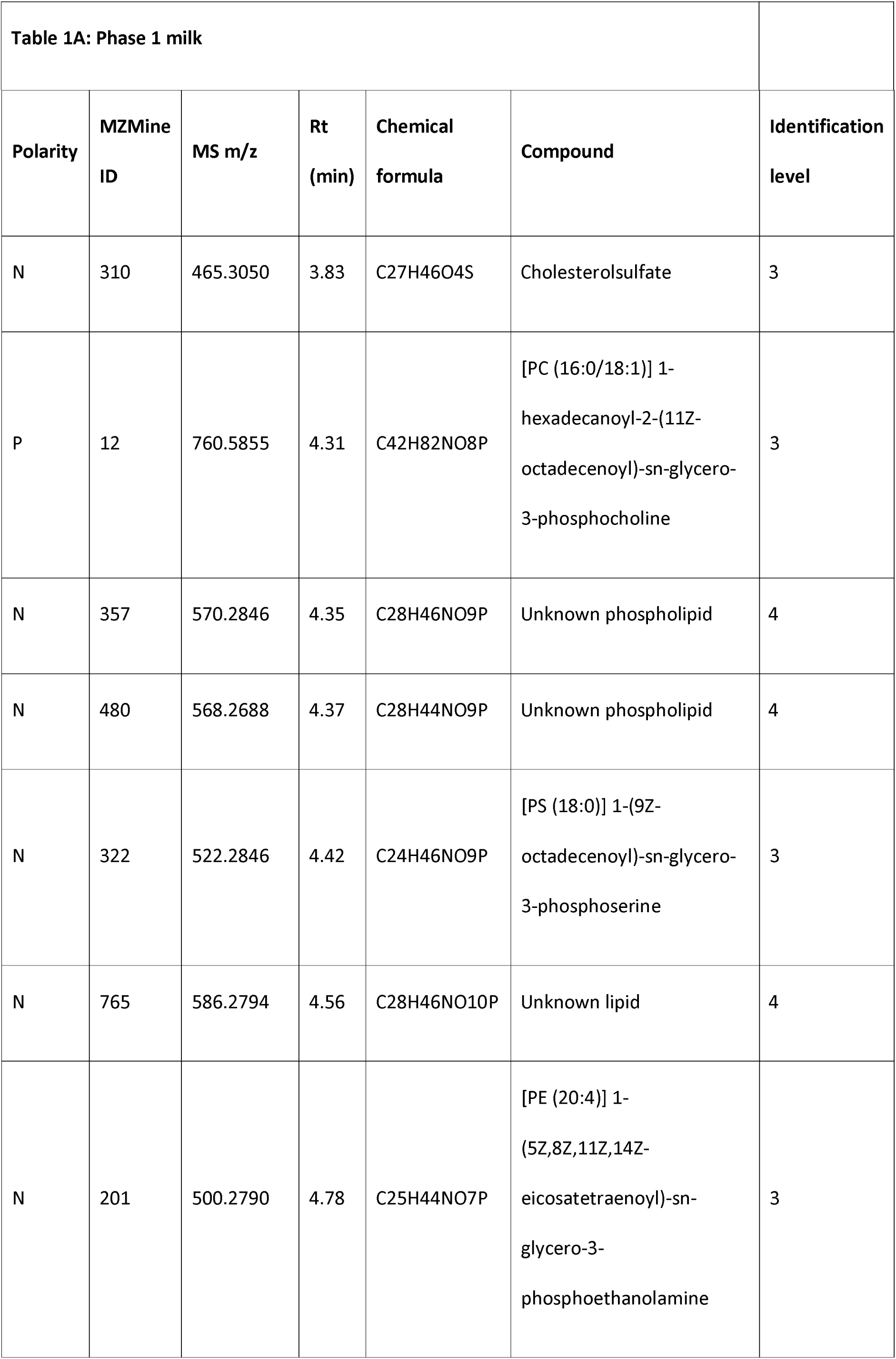
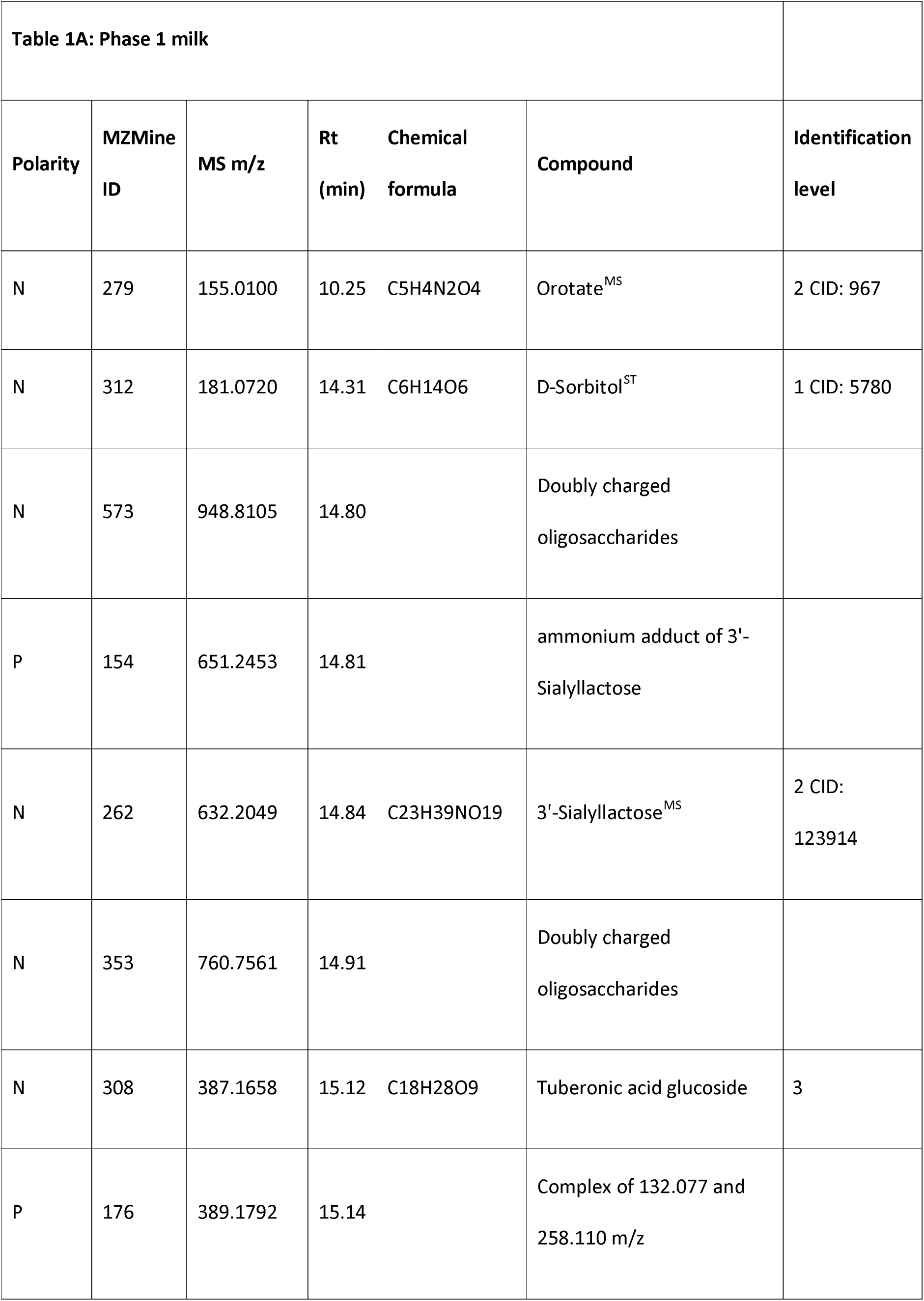
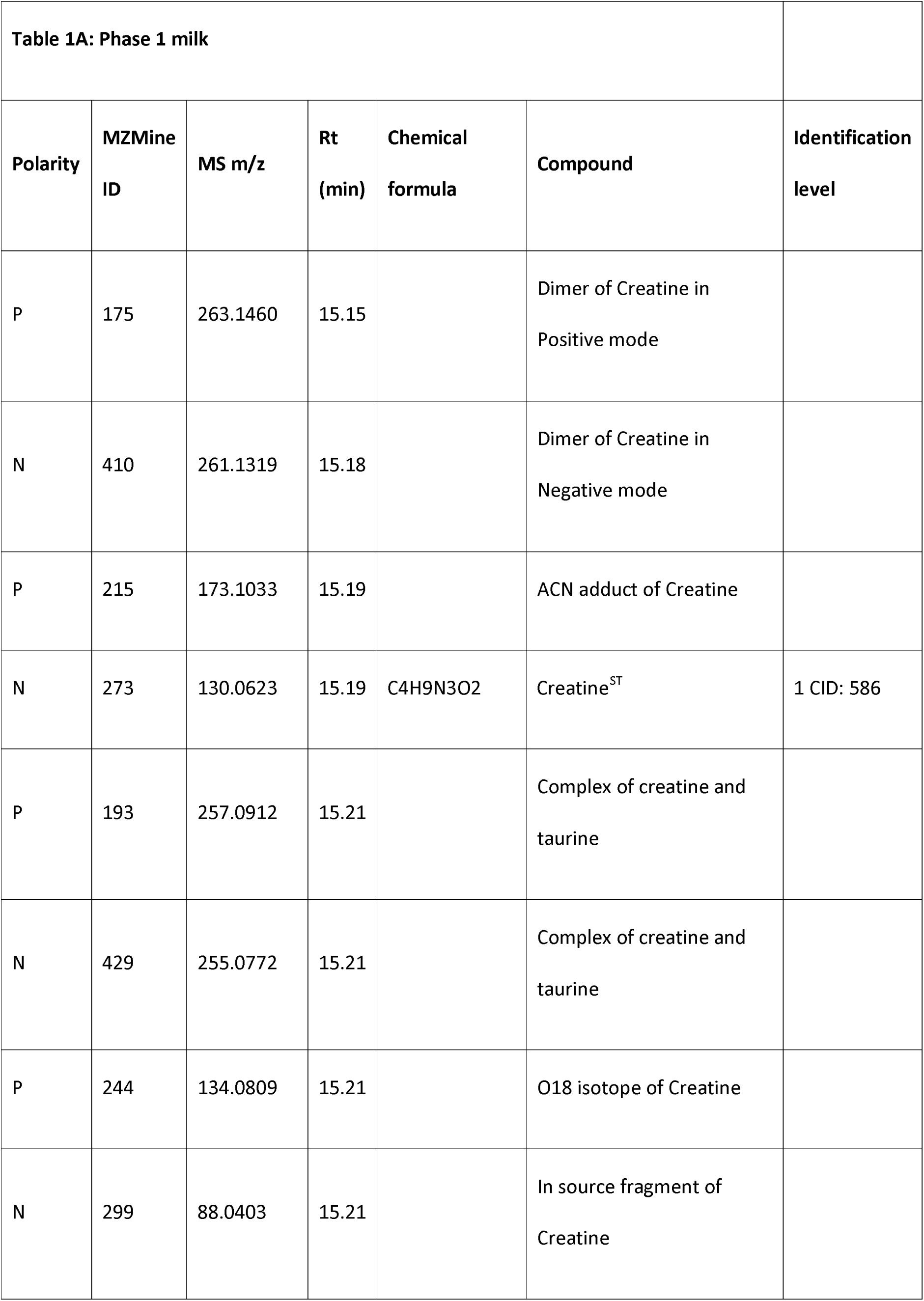
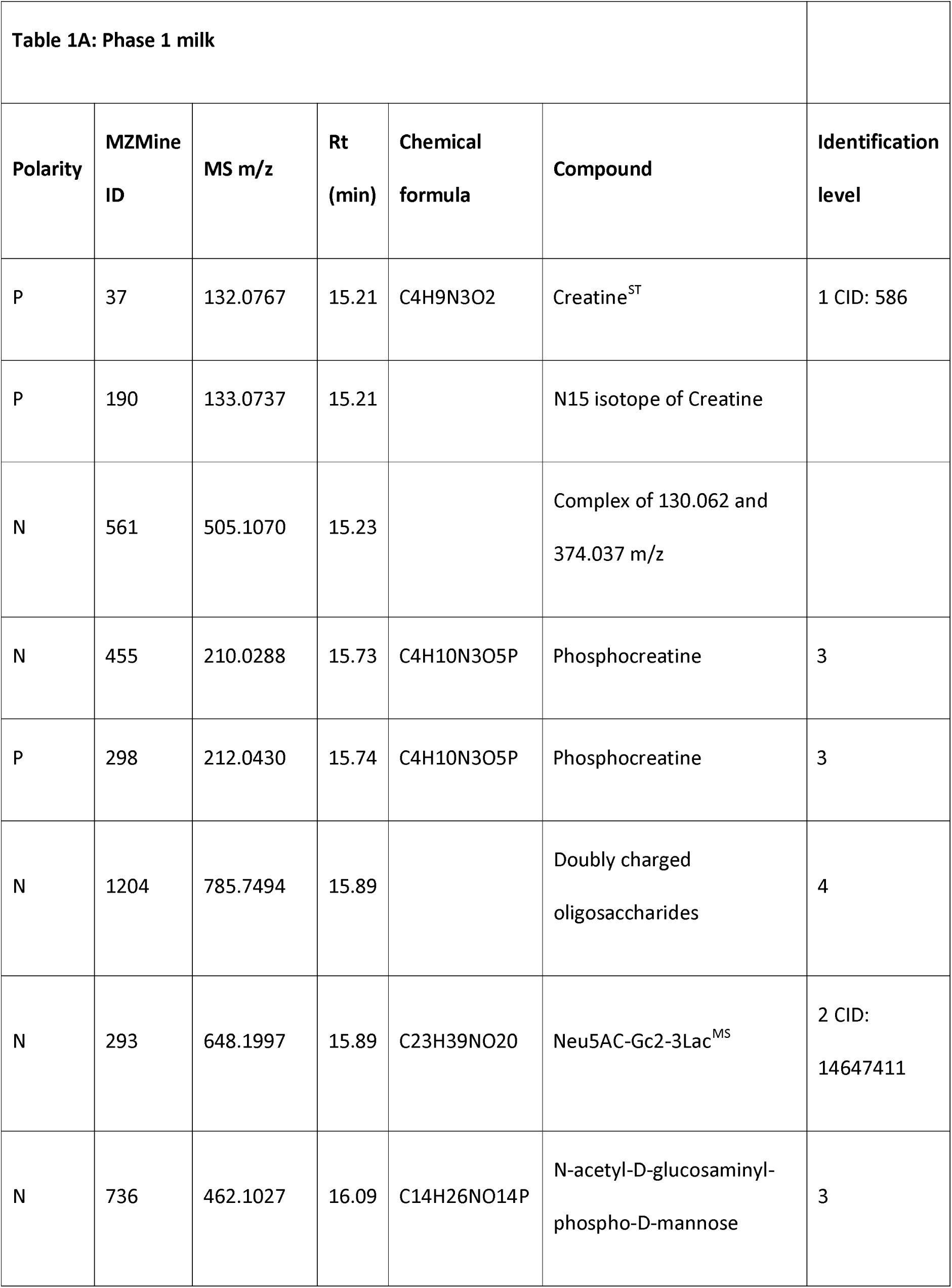
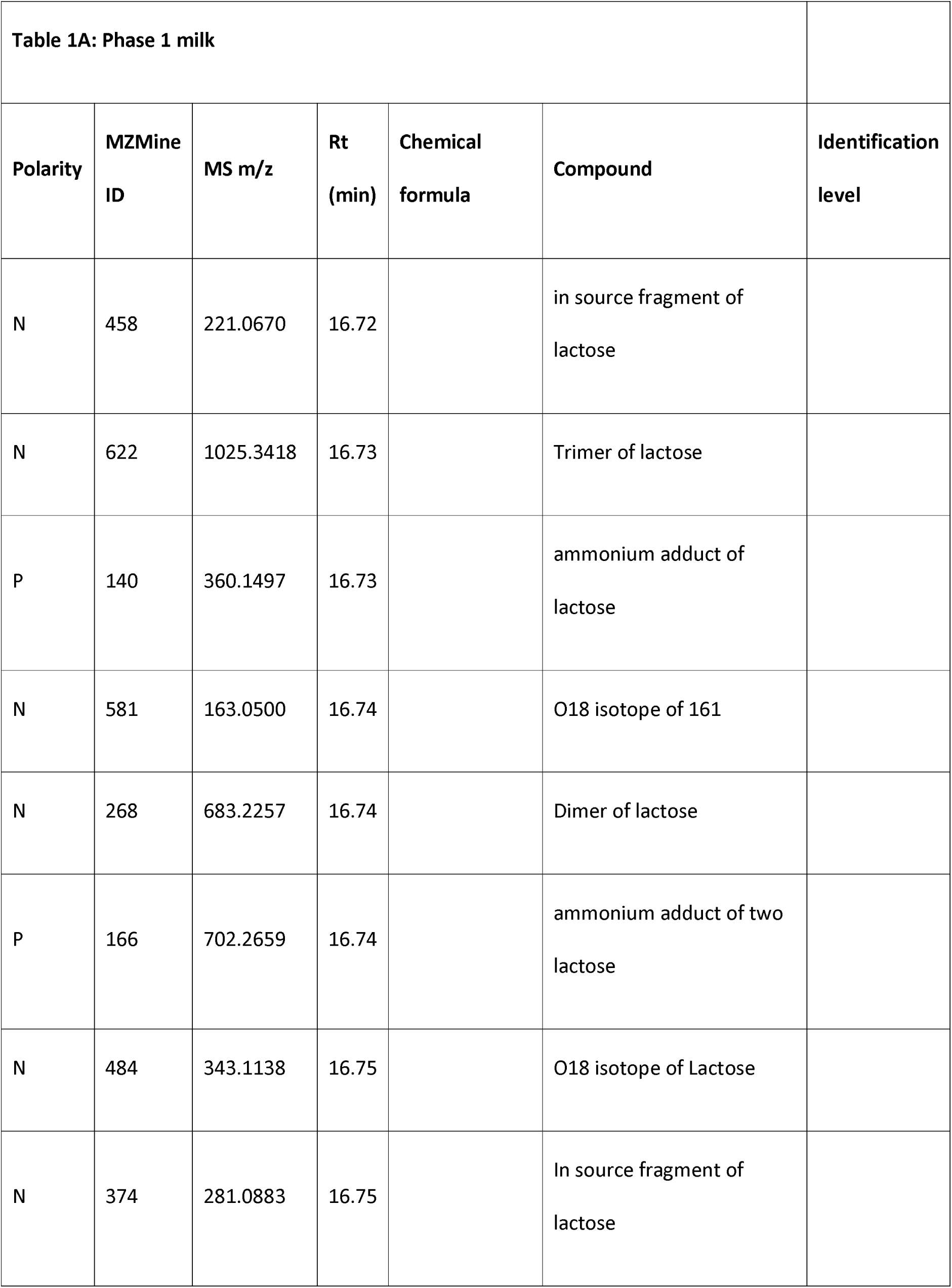
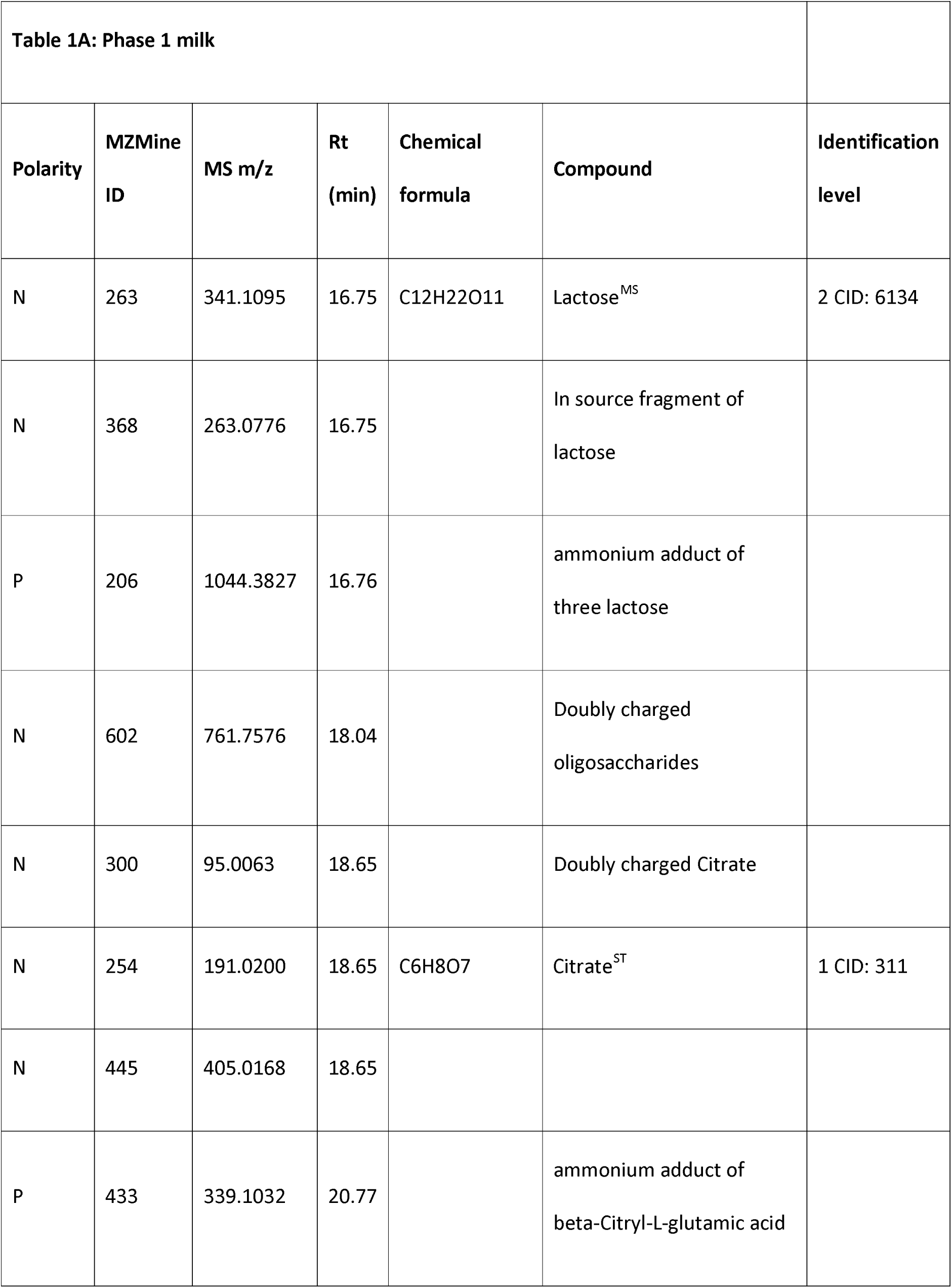
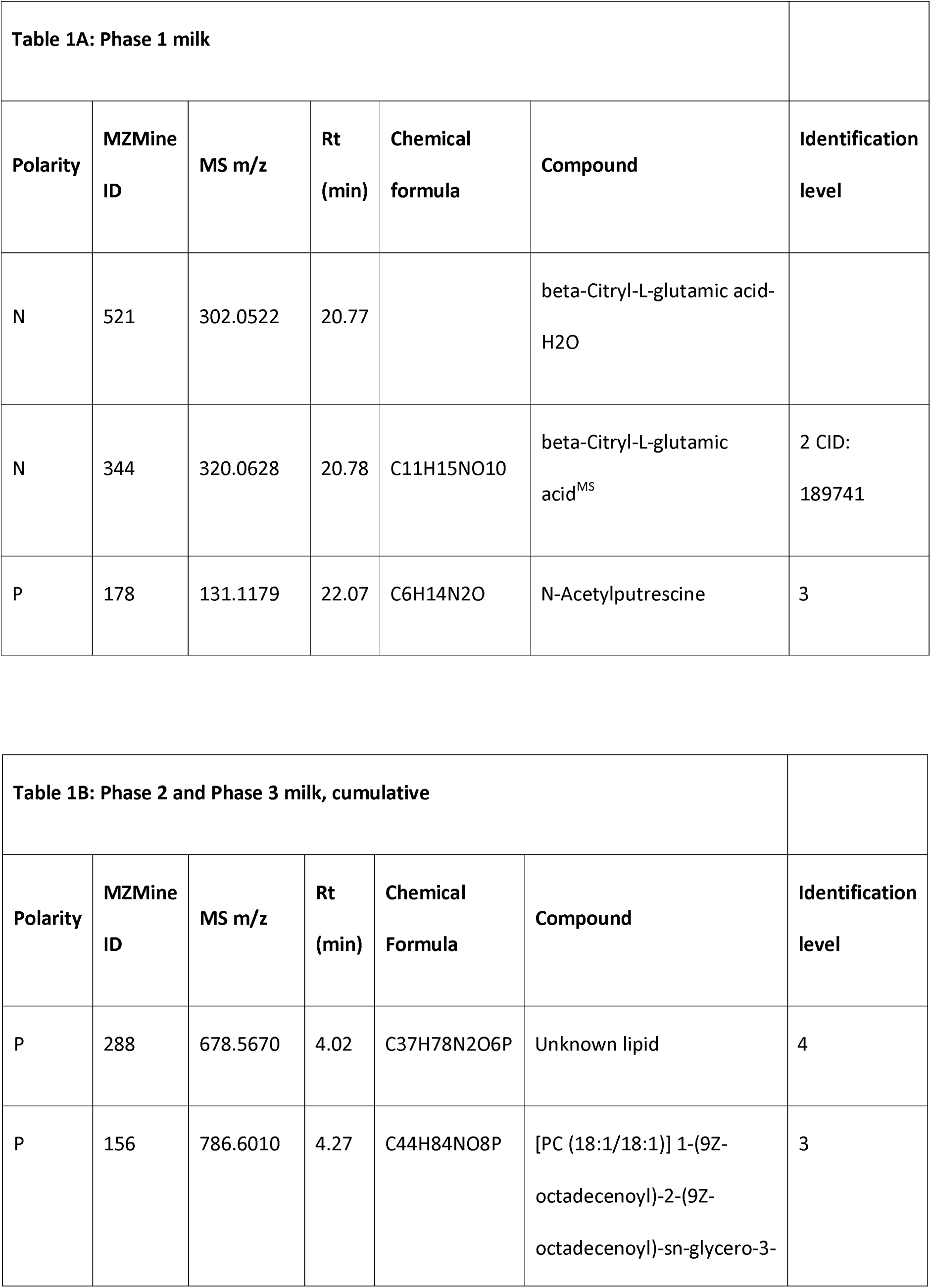
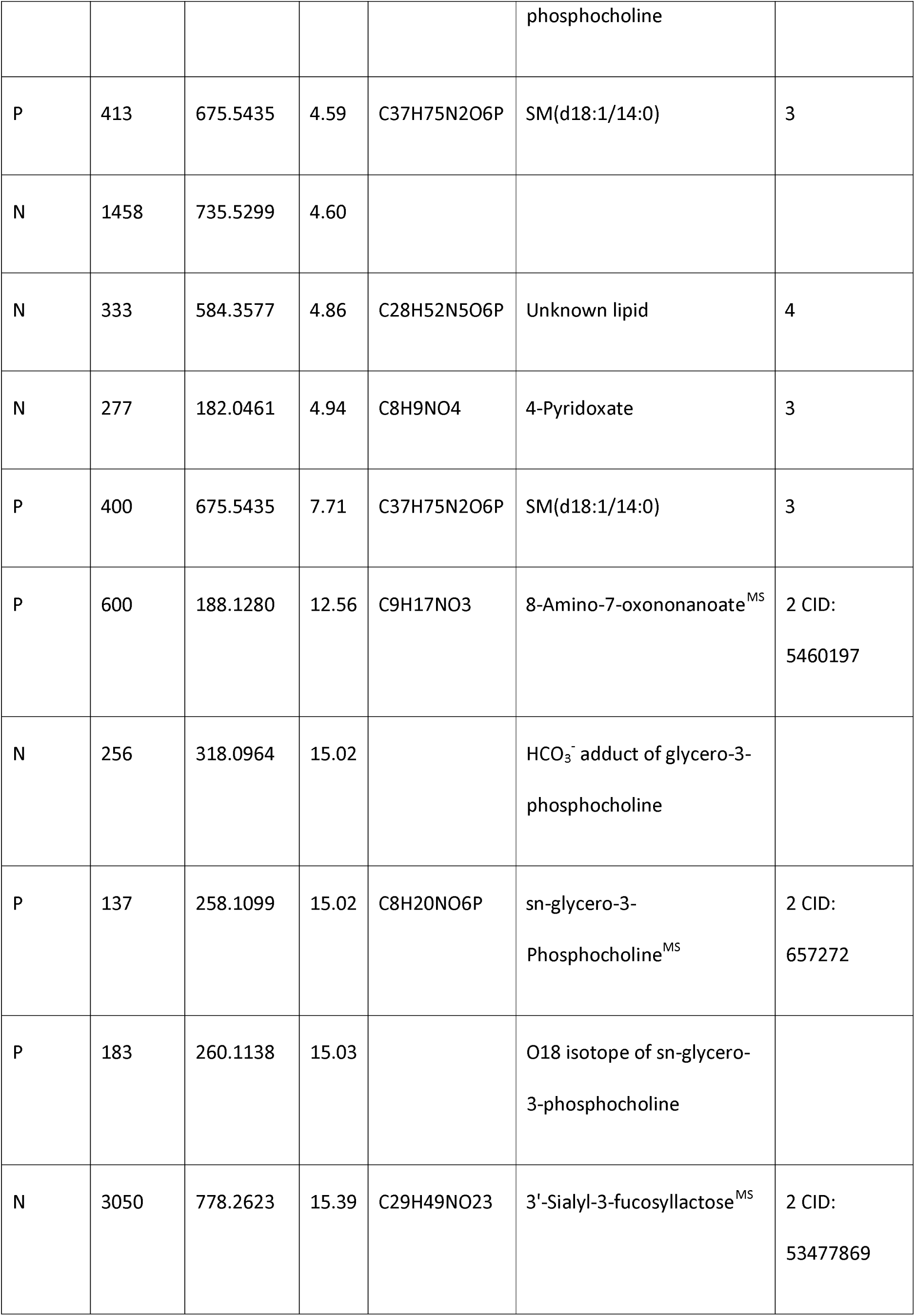
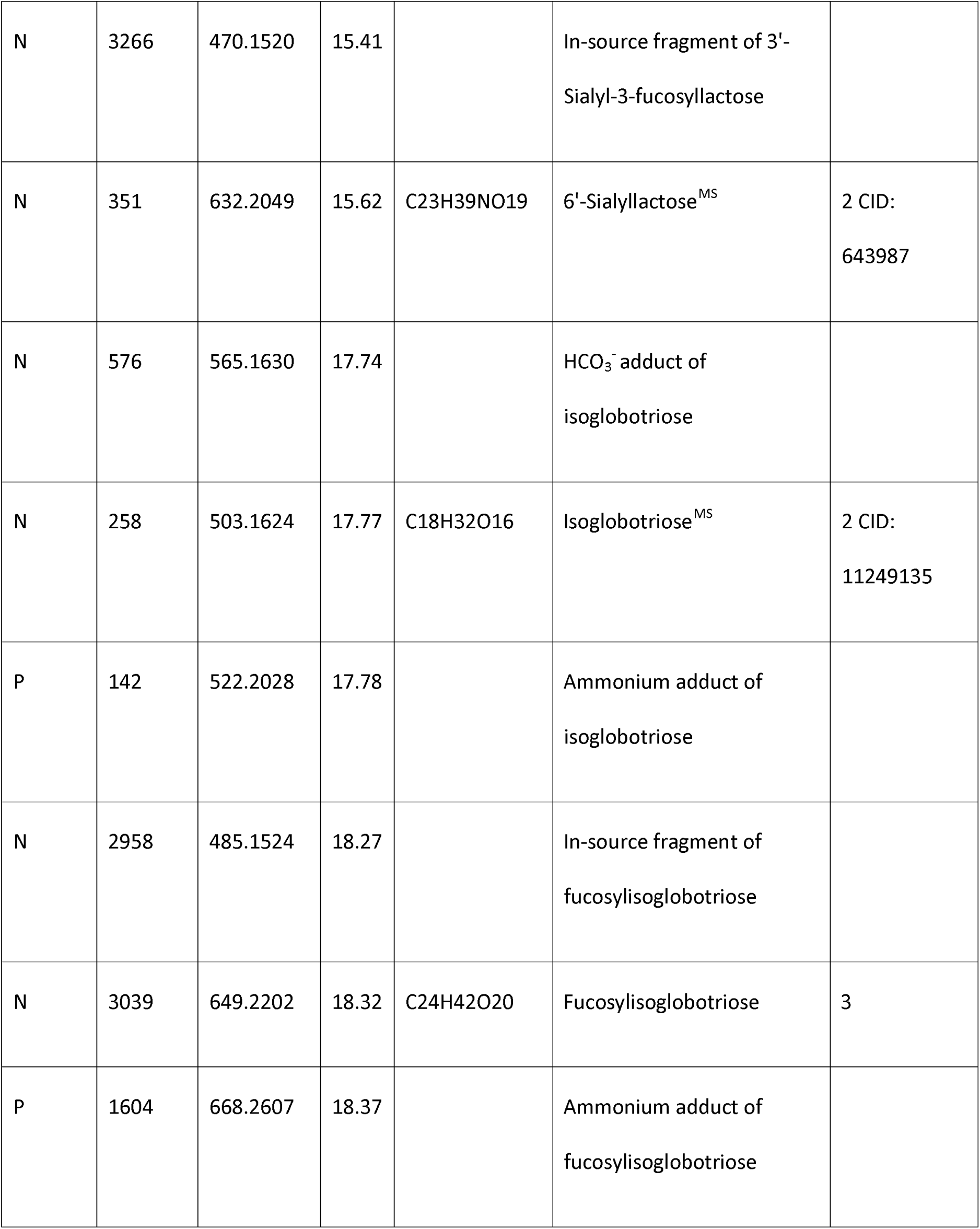
The dominant compounds in giant panda milk in Phase 1 and subsequent phases of early lactation. The 50 most abundant liquid chromatography-high resolution mass spectrometry (LC-HRMS) signals in Phase 1 (the first week of lactation; Table 1A), and the 20 most abundant in Phase 2 and Phase 3 together (7 days and after; Table 1B) milks. Relative abundances were estimated from areas under LC peaks. The compounds are listed in the order of their LC retention times (Rt). Italicised names indicate non-proton adducts and complex ions identified by MZMine 2.10, and confirmed by manually checking the raw LC-HRMS data. The metabolite annotations are based on the Metabolomics Standards Initiative (MSI) identification levels; level 1, retention times matched with authentic standards (labeled as ST); level 2: identified by MS/MS (labeled as MS); level 3: accurate mass; and level 4 unknown. The metabolites identified at levels 1 and 2 were also labeled with the compound identifiers (CID) codes from the PubChem database.

The LC-HRMS features identified and shown in Table 1 represent a diverse array of chemical classifications, and include amino acids, oligosaccharides and lipids. Because milk contains many compounds in addition to those of strictly nutritional value for nursing offspring, it was not unexpected to observe compounds that may reflect physiological or metabolic processes in the mother, the products of which may percolate into milk, as well as the changing needs of the developing altricial infant.

It has been reported that lactose levels are relatively low in milks of pandas and many members of the Carnivora, compared with milks of other species such as bovids and humans (17, 38-41). The data presented here for giant pandas indicate that lactose levels are low and decrease with time after birth (Fig S3).

Two sialylated disaccharide isomers exhibited inverse time-dependent trends (compounds N262 and N351; see insets Fig 2). Their MS/MS spectra displayed common fragments but in different ratios (Fig S4.1). By comparing the fragmentation patterns with published data (42), these two compounds were confirmed as 3’-sialyllactose (N262, high in the first week of lactation) and 6’-sialyllactose (N351, low in the first week of lactation). Differences in these two disaccharide isomers with time after birth have been noted in the milk of other species, though with different relative kinetics from that which we observed for giant pandas (43), or with different relative concentrations between individuals (44). Another important sialylated disaccharide identified here was putatively identified as Gc2-3Lac, which lacks an oxygen atom in the sialic acid residue in comparison to 3’-sialyllactose (Fig S4.2). Sialyllactoses and other sialylated oligosaccharides recently have been reported to have potential benefits in promoting resistance to pathogens, maturation of the gut microbiome, immune function and cognitive development (3, 45-47). Levels of these sialylated oligosaccharides were notably low in all three artificial milk formulae tested in this study. This observation and other differences of potential physiological significance between maternal and artificial milk are discussed below.

OPLS-DA was performed on the post-20-day samples of YY and XYT (Fig S5.1; see also Figs S5.2 and S5.3 for statistical validation) in order to describe the key components responsible for the discrimination between the two individuals. Based on the magnitude of their contributions to the model, the top 20 LC-HRMS features were extracted for each panda, and are listed in Table S4. Most of these are low molecular weight metabolites rather than the larger components such as lipids and oligosaccharides. For instance, a significant difference in the levels of several acylcarnitines might indicate differences in fatty acid metabolism between these two individuals. For the later samples, maternal factors such as age, genetics, epigenetics, diet, disease history, and parity may influence the composition of milks. This emphasises the importance of maternal health, stress and physiology on milk composition and, in turn, the impact of maternal well-being on infant development.

The low sample number precluded analysis of the potential effects of age, genetics and parity in this study. However, the influence of maternal diet and medical history on milk composition would have been an important evaluation to subtract from the metabolomic changes that reflect neonatal needs. For example, levels of phenol sulphate in the milk of giant panda mothers were initially very high, and then declined steadily until their disappearance at ca. 30 days. The pattern corresponds with the 7-14 day period of anorexia that is typical of giant panda mothers in the immediate post-parturient period. Data necessary for correlating nutritional and maternal medical features with metabolites in the milk were not available for this study, but would be important to include in future studies of bear milks. This will be particularly important in the comparison of the milk metabolome among species of bears that hibernate during parturition and early lactation (black bears, brown bears, polar bears) and those that do not (giant pandas, sloth, sun, and Andean bears).

Compounds may increase or decrease with time in milks for several reasons. The simplest explanation may be that they are directly nutritive and their abundances change according to the developmental needs of the neonate. Another is that they may be involved in innate immunity by, for example, acting in the blockade of cellular receptors against bacterial pathogens, as is seemingly the case for certain oligosaccharides (45). Also, as is thought to be the case in humans (45), some oligosaccharides may neither be digested nor involved in anti-bacterial defence, but instead act to control the establishment of an appropriate microbiome.

### Potential of metabolomics statistical methods in inferring associations between the changing milk metabolome and cub growth rate

Some compounds showed clear increases or decreases in abundance with time and progression through the three phases of early lactation. Deriving cause and effect associations between changes in metabolite abundance and cub growth rates would be speculative at this stage, especially given that the growth rates of the cubs for which we have information are a composite of nutrition from both natural nursing and supplementation with artificial formula. Nevertheless, in order to begin to explore potential associations, we applied OPLS regression analysis to link the LC-HRMS features that change in relative abundance (X-variables) to the rate of panda cub weight gain (Y-variables; Fig 3). For this, we were confined to the one panda mother from whom milk had been sampled over a sufficient period of time and her cub (YY and cub; Fig 3; Fig S1). The change in growth rate of this cub is given in Fig 3B. It shows the weight drop immediately after birth that is normal in mammals, followed by a rapid increase which peaked at around 7 days after birth, and then slowed but remained positive.

**Fig 3.**
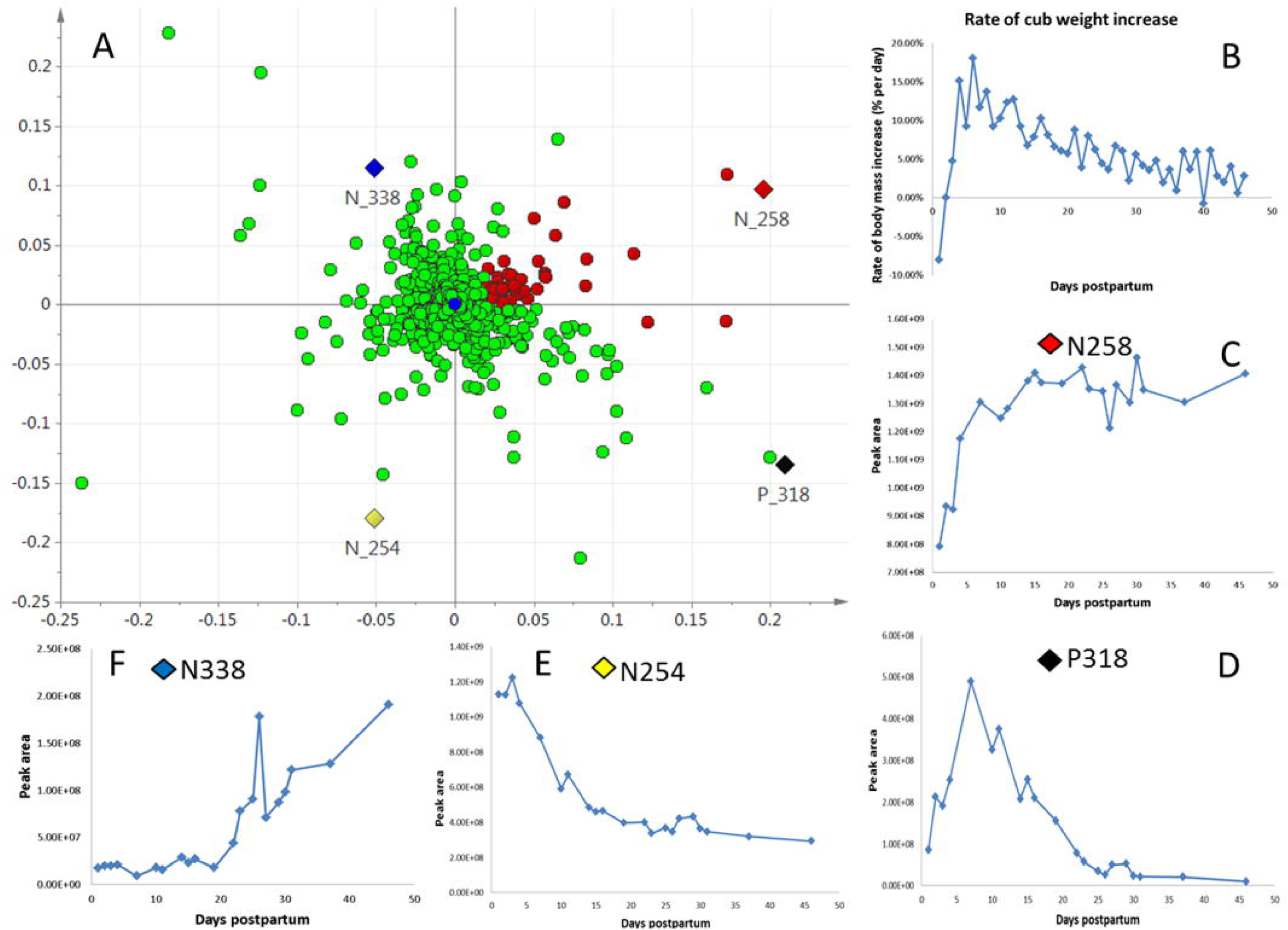
Compounds in giant panda milk tentatively correlated with cub growth rate. (A) OPLS loading plot for 21 serially collected samples from giant panda Yuan Yuan (YY). Components selected for testing for correlation with rate of cub body weight change are highlighted in red; see main text for selection criteria. X-variables, Pareto scaled LC-HRMS data; Y-variable, rate of body weight change over time of YY’s cub. (B) Body mass changes for YY’s cub over the milk sampling period (see also Fig. S1). (C, D, E, F) Changes in abundance with time of the compounds indicated with diamonds in (A) selected to illustrate diverse correlations with weight gain by the cub. Compound N258 showed positive correlation, whereas compound N254 showed negative correlation, and compound P318 exhibited a more complex pattern. Y-axes represent relative abundances of each compound as estimated from areas under peaks calculated from HILIC-HRMS data, x-axes represent time after birth. Putative initial identifications were N258, isoglobotriose; P318, methyl-imidazole acetate; N254, citrate; N338, 3-methyl-2-oxopentanate.

An OPLS regression analysis was used statistically to link the LC-HRMS features to the rate of the cub’s weight gain; the permutation test is shown in Fig S6. The loading plot is shown in Fig 3A. Compounds that were further identified as being of potential interest are indicated in red, and were selected as follows. Theoretically, highly-correlated X-variables have a high positive correlation score in OPLS regression analysis, with a zero-approaching orthogonal score located at the right of the horizontal axis in the loading plot. Four line graphs are displayed (Fig 3C-F), illustrating notable examples of such correlations. The change in relative abundance of component N258 (isoglobotriose; Fig 3C) follows a roughly similar profile to the rate of the cub’s weight gain (Fig 3B). In contrast, components located in the other three quadrants of the plot (highlighted in the loading plot, and their changes with time shown in Figs 3D-F) show different associations with growth, one in particular showing a negative association (N254; citrate, possibly an excretory product rather than a nutrient; Fig 3E). In order to focus further on compounds that increase along with body weight, we selected by visual inspection 50 of the most abundant compounds based on relative intensity (Fig 3A, red dots) appearing in the x- and y-axis positive quadrant, and a few abundant components falling slightly y-negative to allow for slight errors.

Further identification of these 50 components was carried out by comparison of LC-MS retention time with standards and/or by MS/MS analysis (Table 2). Compound N258, the relative abundance of which showed a positive correlation with the cub’s rate of weight gain, was identified as isoglobotriose. The change in relative abundance of this trisaccharide over time showed a similar pattern among all three pandas (Fig S7). It was low initially, and increased to a plateau at about 15 days. Isoglobotriose dominates in ursid milks [34], but is notably low in the artificial formulae that we examined, lower even than the level at the initiation of our lactation sampling series (Fig S7).

**Table 2.**
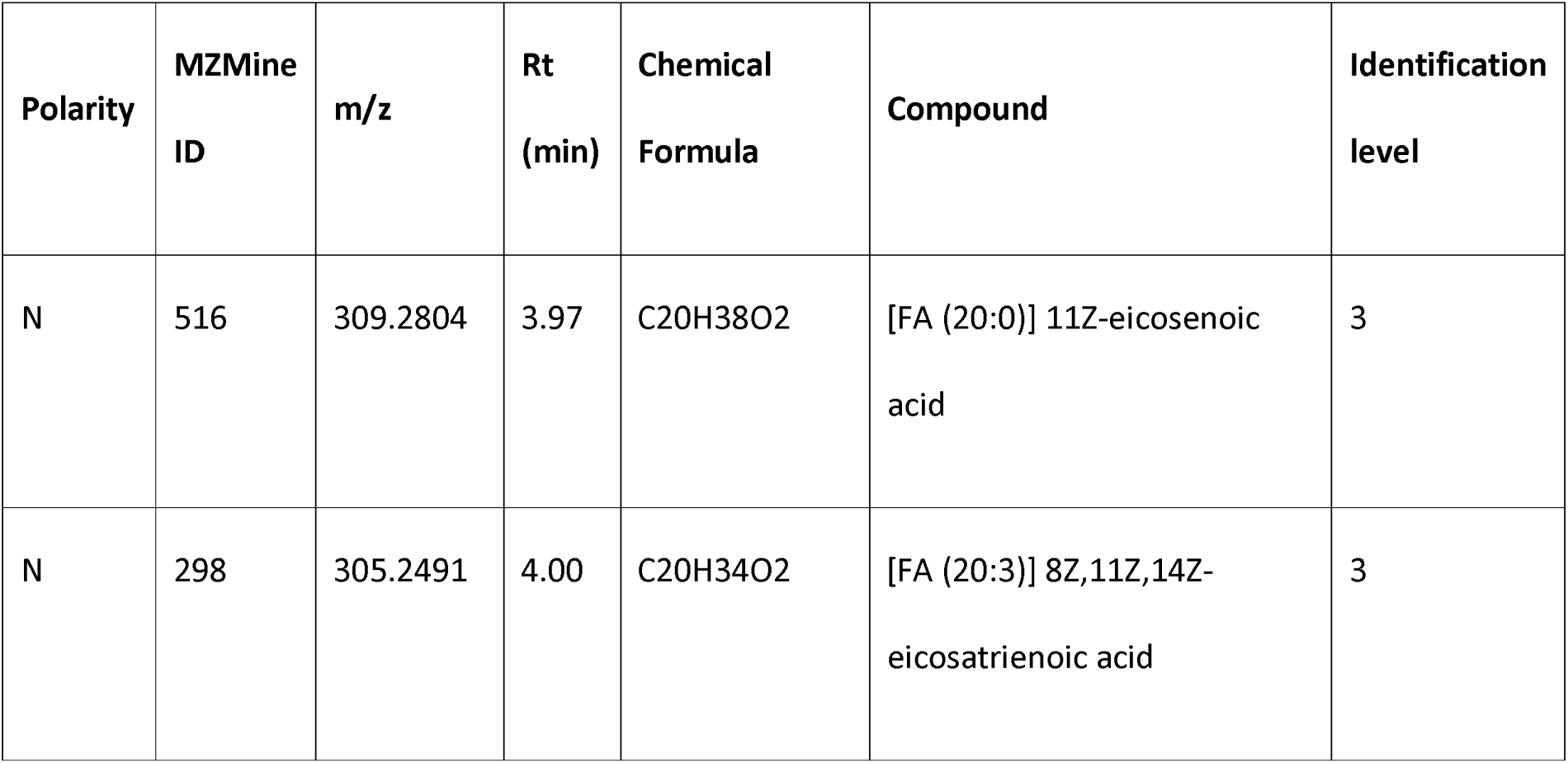
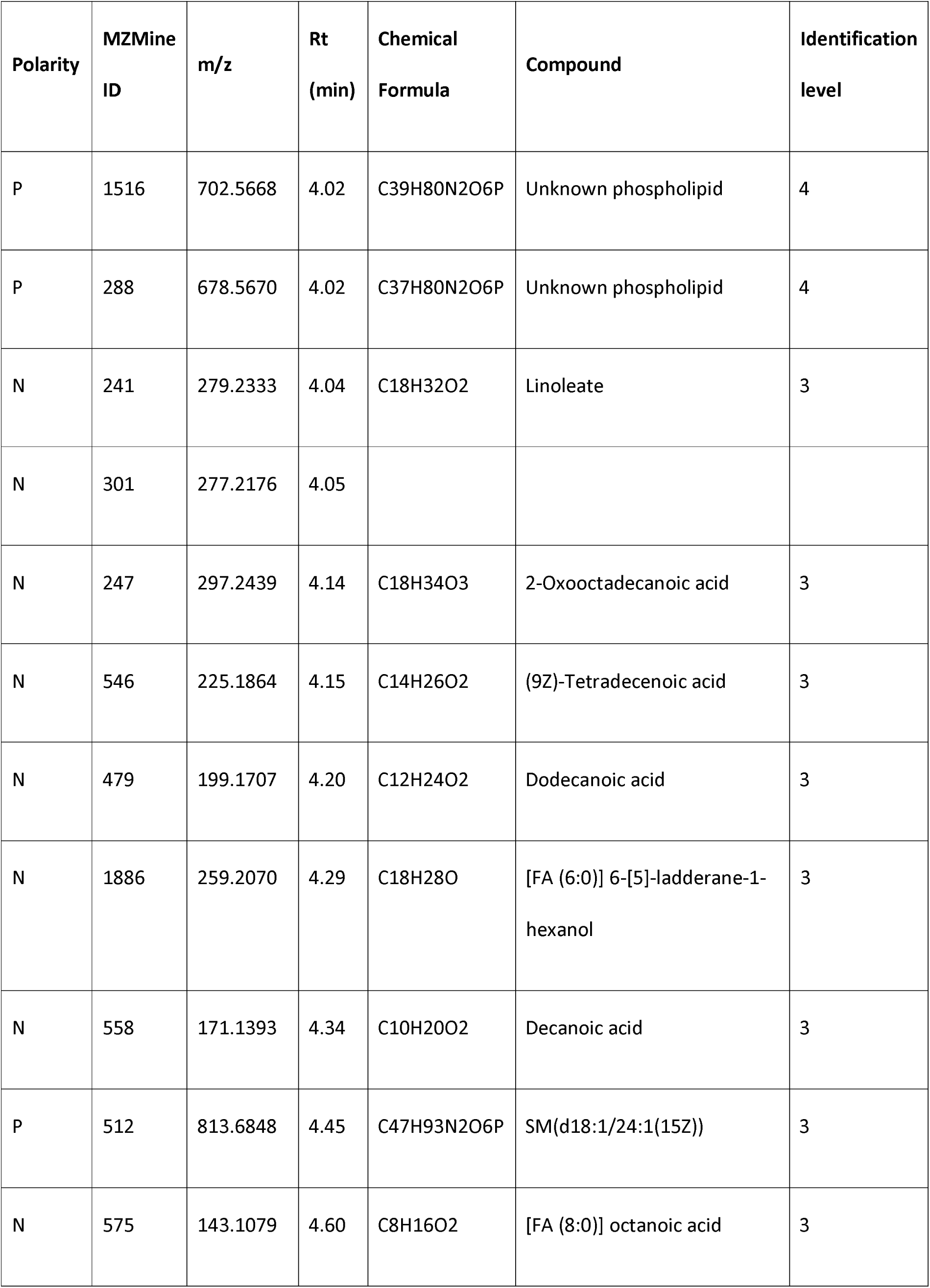
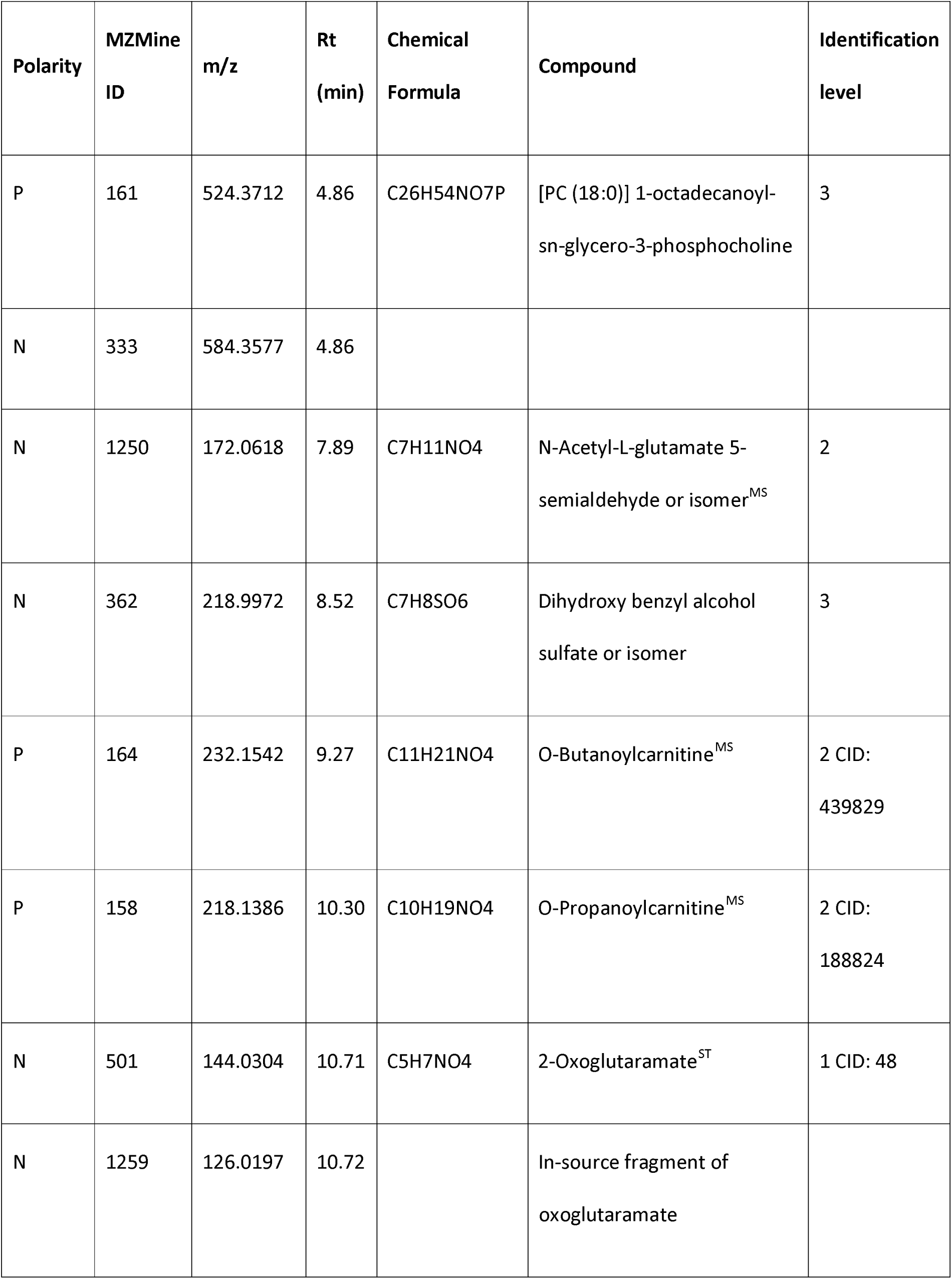
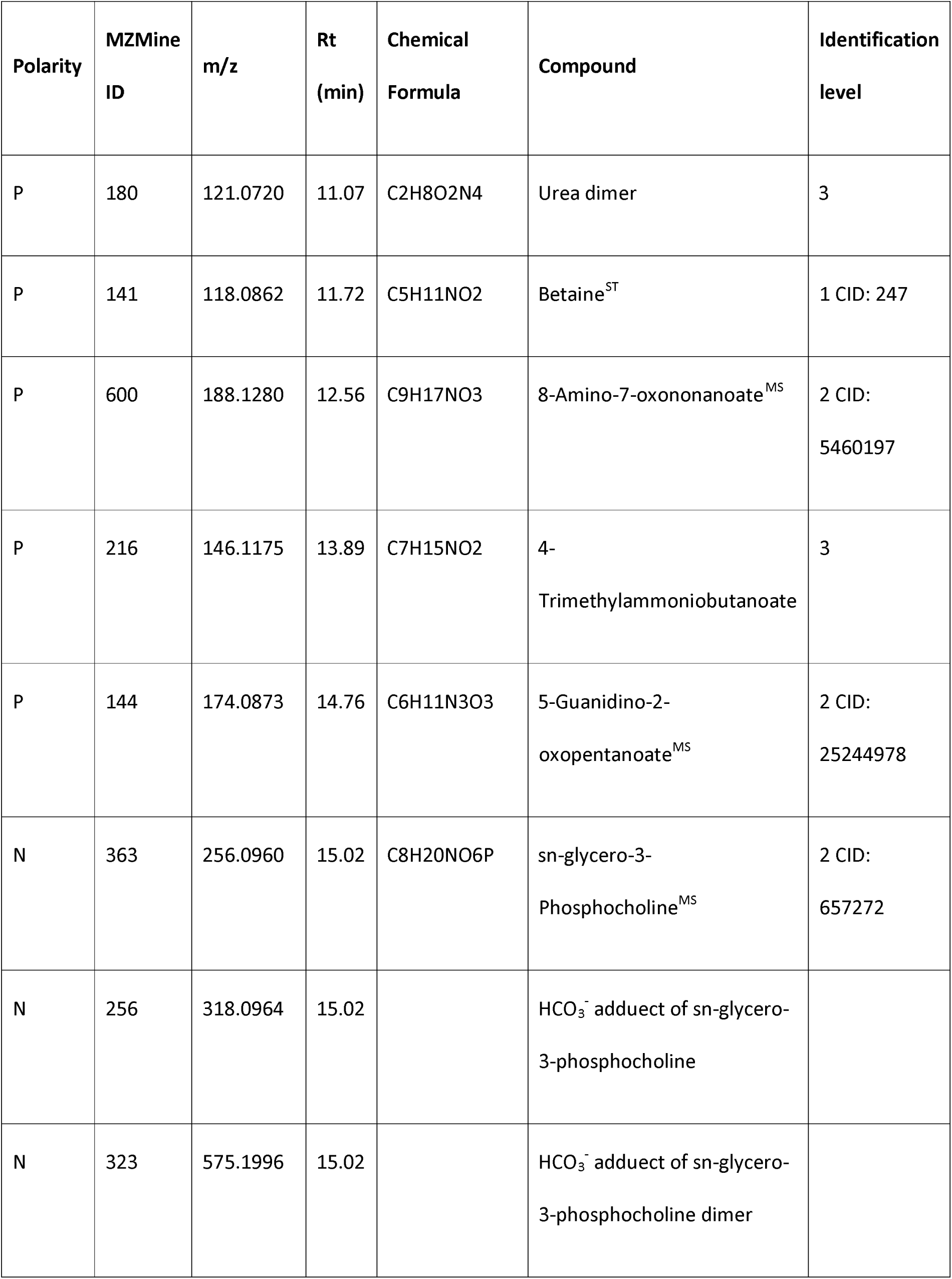
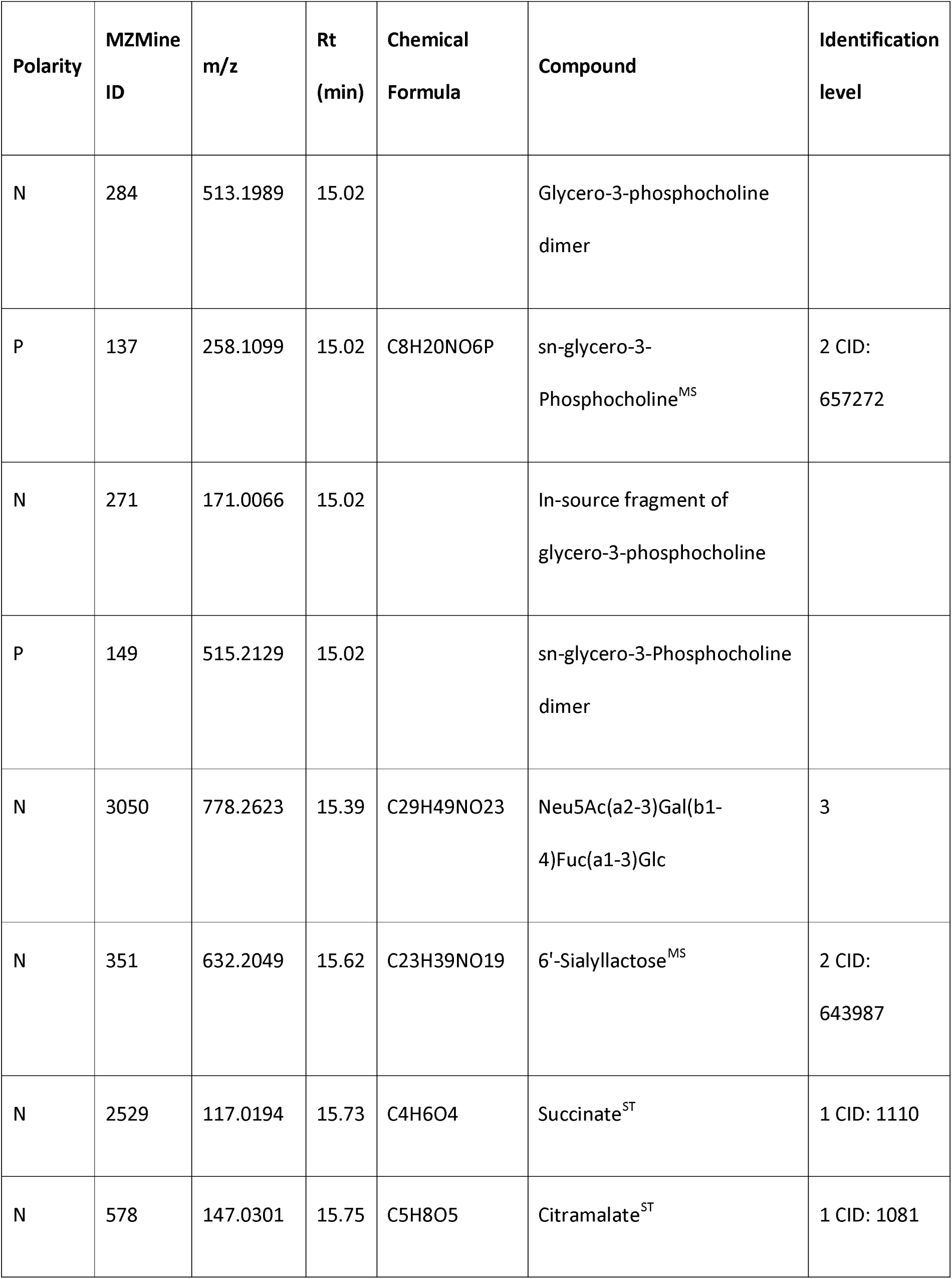
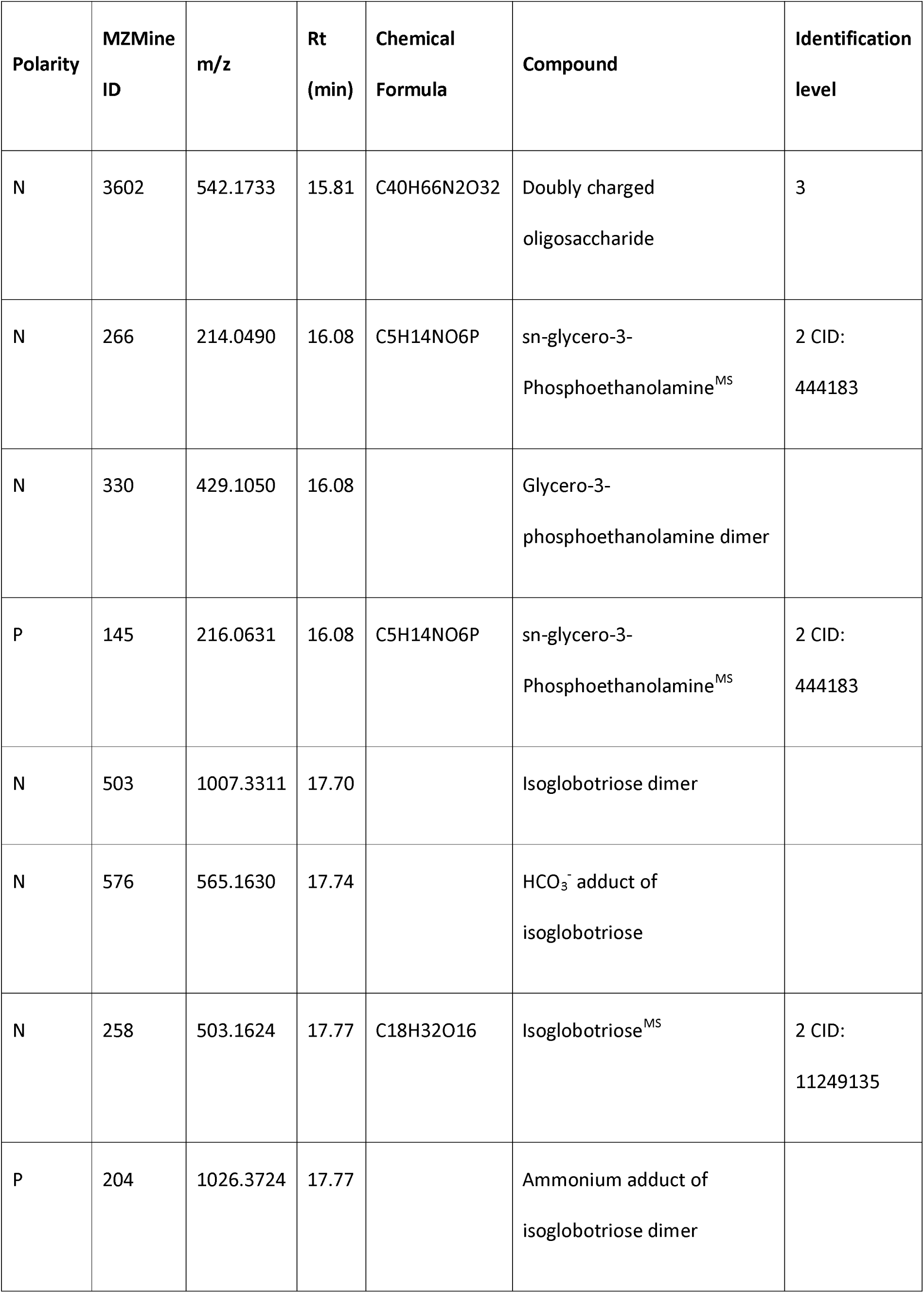
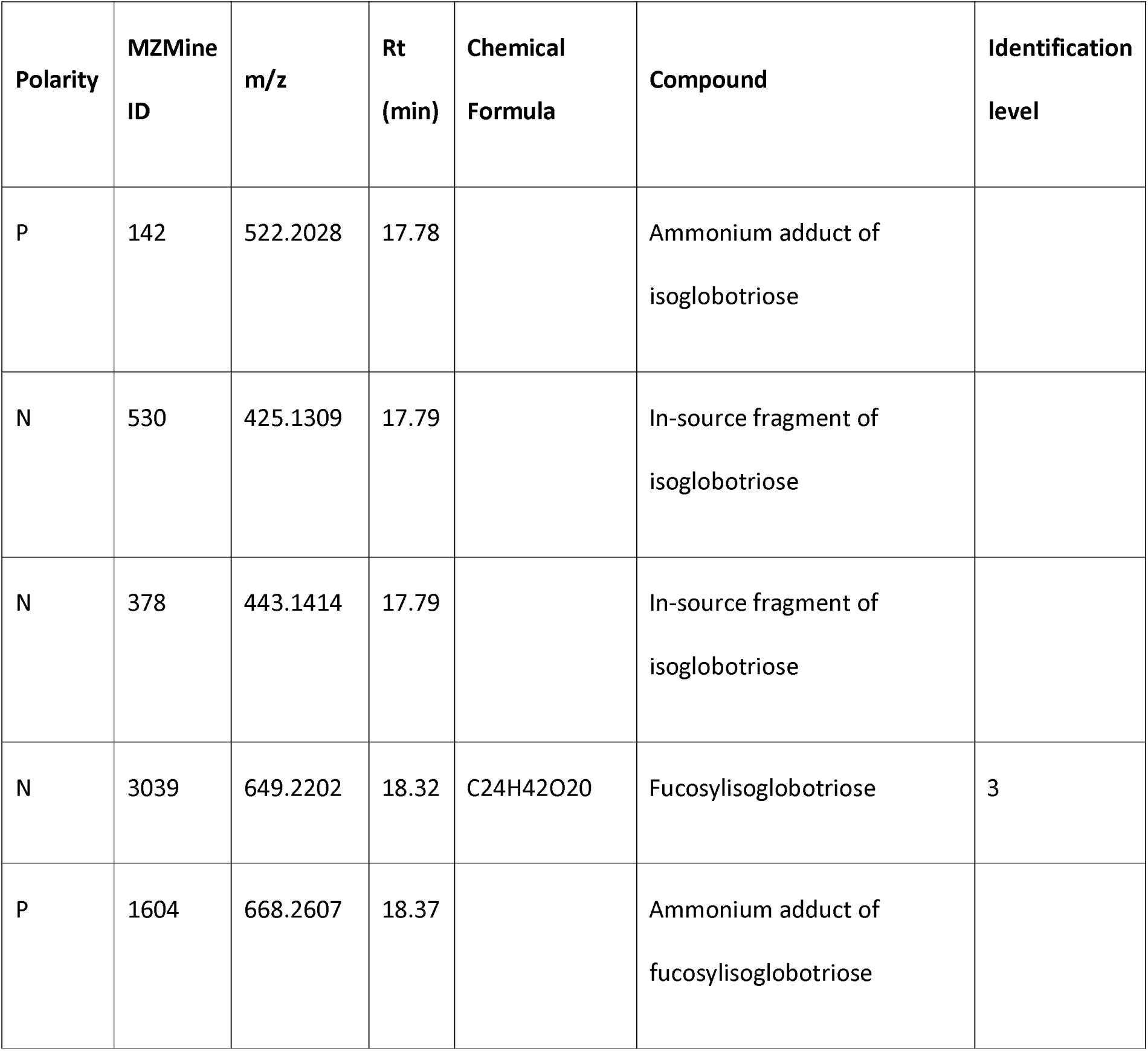
Compounds in giant panda milk tentatively correlated with cub growth rate. The 50 most abundant LC-HRMS signals selected from the OPLS loading plot of of YY’s 21 samples. The LC-MS features are listed in the order of their LC retention times (Rt). Italicised names indicate non-proton adducts and complex ions identified by MZMine 2.10, and confirmed by manually checking the raw LC-HRMS data. The metabolite annotations are based on the Metabolomics Standards Initiative (MSI) identification levels. Level 1, retention times matched with authentic standards (labelled as ST); level 2, identified by MS/MS (labelled as MS); level 3, accurate mass; and level 4, unidentified. The metabolites identified at levels 1 and 2 are also labelled with the compound identifiers (CID) codes from the PubChem database.

Artificial formulae are commonly based on bovine milk, which, in contrast to bear milks, has a low content of oligosaccharides and high lactose (48). In this study of giant panda milks, we found that lactose peaked alongside isoglobotriose initially, and then fell to a plateau at about 18 days (Fig S3). These observations support the clinical observations that bear cubs are not adapted to digest lactose-rich formulae (49, 50), as is also known to be the case for some domestic Carnivora, such as cats (51).

While time-resolved lipidomics of panda milk remains to be carried out, it bears mention that n-glycero-3-phosphocholine and sn-glycero-3-phosphethanolamine were identified here to follow a pattern similar to that of isoglobotriose in relation to cub growth rate (Figs S8 and S9). These compounds provide the hydrophilic head groups of many phospholipids involved in cell membrane construction, and are also precursors of glycerol-3-phosphate, which is important in many biosynthetic reactions. Fatty acids that were identified included linoleic acid, which is a nutritionally-essential polyunsaturated omega-6 fatty acid (Table 2).

### Concluding remarks

This study describes the application of an HILIC-HRMS method and data processing strategy to the metabolomic analysis of 55 giant panda milk samples collected from three mothers during the first two months of lactation. In spite of the fact that the current study is based on only three animals, this is the most comprehensive study of ursine (or any Carnivoran) milks reported to date with regard to the number of samples and, in particular, serial coverage of the lactation period from birth. The PCA score plot indicated changes in milk components and differences that developed among individual mothers over time. Three phases of early lactation were identified in this analysis, demarcating the first week and then after the third week of lactation. Application of metabolomic methods such as ours to the milks of species with different lactation strategies and physiologies, and with similarly or even less-altricial new-borns, will enhance our understanding of the diversity of mammalian infant support in the immediate postpartum period.

With regard to bears, this study illuminates unusual features of ursine lactation as may be adapted to provide for their unusually altricial neonates. Our findings highlight the intricacies of maternal lactation physiology and the chronologically dynamic and complex balance of nutritional and immunological features, and other components that meet the changing requirements for infant ontogeny after birth.

Of particular note, our analyses emphasise the inadequacy of artificial milk replacers for bear cubs, and the challenges in formulating such replacers with any fidelity and without danger of doing harm to growing cubs. Over-supplementation with a specific component, or otherwise disrupting the balance with discriminatory supplementation with, for example, oligosaccharides, could be deleterious to cub health and development. Our findings on statistical correlations between compounds in giant panda milks and the rate of cub growth are illustrative of principle and cannot be taken as definitive. The methods we developed and tested will only be useful in further studies in which, for instance, food supplements have not been given to mothers or cubs, and more natural feeding regimes have been allowed. Artificial formulae are clearly necessary for the rescue of orphaned cubs, but, where lactating mothers are available, they are not warranted and could be deleterious even if given as a supplement to mothers’ milks. Moreover, milk is but one vital component of maternal care for the development of a physically and behaviourally healthy cub, and it is critical to allow the mother-cub bond to proceed its natural course with as little human intervention as possible.

## Acknowledgments

We are grateful to the staff of the Chengdu Research Base of Giant Panda Breeding for providing the milk samples.

## Supporting information

**Table S1.** Giant panda mother information and dates of milk sample collection.

**Table S2.** MS data of the significant components in giant panda milk selected by multivariate statistical analysis.

**Table S3.** LC-HRMS features verified by comparison with the retention times of authentic standards.

**Table S4.** Compounds detected in the milk samples of giant pandas YY and XYT 20 days postpartum, and which differentiated the two individuals.

**Fig S1.** Body weights of three giant panda cubs over their first 60 days after birth.

**Fig S2.1.** OPLS-DA score plot of 55 giant panda milk samples before and after 7 days of lactation.

**Fig S2.2.** Statistical validation of the OPLS-DA model by permutation analysis.

**Fig S3.** Relative abundance of lactose in the milk samples of the three giant pandas with time postpartum.

**Fig S4.1.** Extracted ion chromatograph of 3’ and 6’-Sialyllactose and their MS/MS spectra.

**Fig S4.2.** Extracted ion chromatograph of Gc2–3Lac and its MS/MS and MS/MS/MS spectra.

**Fig S5.1.** OPLS-DA score plot of YY and XYT milk samples 20 days postpartum.

**Fig S5.2.** Statistical validation of the OPLS-DA model by permutation analysis of data from milk samples from giant pandas YY and XYT 20 days after parturition.

**Fig S5.3.** OPLS-DA S-plot of YY and XYT milk samples 20 days postpartum.

**Fig S6.** Statistical validation of the OPLS-DA model by permutation analysis of data from 21 milk samples from giant panda YY versus body weight changes of her cub with time after birth.

**Fig S7.** Relative abundance of isoglobotriose, a common component in the milk of bears, in the milk samples from the three giant pandas with time postpartum.

**Fig S8.** Relative abundance of sn-glycero-3-phosphocholine in the milk samples of the three giant pandas with time postpartum.

**Fig S9.** Relative abundance of sn-glycero-3-phosphoethanolamine in the milk samples of the three giant pandas with time postpartum.

